# SNP Identity Searching Across Ethnicities, Kinship, and Admixture

**DOI:** 10.1101/574566

**Authors:** Brian S. Helfer, Darrell O. Ricke

## Abstract

Advances in high throughput sequencing (HTS) enables application of single nucleotide polymorphism (SNP) panels for identification and mixture analysis. Large reference sets of characterized individuals with documented relationships, ethnicities, and admixture are not yet available for characterizing the impacts of different ethnicities, kinships, and admixture on identification and mixture search results for relatives and unrelated individuals. Models for the expected results are presented with comparison results on two *in silico* datasets spanning four ethnicities, extended kinship relationships, and also admixture between the four ethnicities.

## Introduction

The United States recently expanded the Combined DNA Index System (CODIS) loci from 13 to 20 in 2016. The current FBI National DNA Index System (NDIS) database has near 18 million profiles (1). Shifting from sizing short tandem repeat (STR) alleles to high throughput sequencing (HTS) will enable resolution of different STR alleles with the same lengths. Illumina recently introduced the ForenSeq (2) sequencing panel that includes SNPs in addition to STRs. This expanded panel of STR and SNP profiles provides additional sample profile information for inferring biogeographic ancestry (BGA) and phenotype (externally visible traits – EVTs). Others have pioneered using SNPs for BGA prediction (3, 4), EVTs (5), mixture analysis (6), and more. Future panels will scale to thousands and tens of thousands of loci with loci selected for multiple forensics insights (6).

Previous work has examined the use of specific SNP panels for identifying populations of individuals (7, 8). This work relies on a specific assay for identifying the ancestry of an individual. This creates a limitation in the ability to apply this work to forensics applications, where SNP markers may not have been genotyped, and would impose a burden of having to re-genotype the DNA with a new panel as the assays are revised. More recently, researchers have been looking to create more generalizable algorithms that can use SNP data to predict biogeographic ancestry and descent (9–15). In this work, a generalizable framework for identifying differences between ethnicities and family members based on a subset of the available SNP data is presented.

## Methods

### Database Generation

SNPs were selected from the Allele Frequency Database (ALFRED) (17), containing four ethnicities with large numbers of well-characterized SNPs: African Americans, Estonians, Koreans, and Palestinians. A total of 39,108 SNPs were characterized across each of these ethnic groups. This information was used to create four initial *in silico* populations of pure ancestry, each containing 15,000 individuals. When multiple sources were listed for a single locus, the median minor allele frequency was assigned to the SNP. The gender of each individual was assigned such that

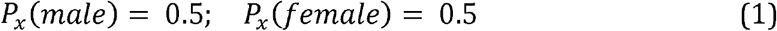

the probability that individual x is male is 0.5, and the probability that individual x is female is also 0.5. Minor allele information was assigned using the allele frequencies provided by ALFRED. Eight more generations were created using these four ethnic populations. Additionally, four populations of the same ethnic groups were created with a different seeding of the random number generator. This results in a starting generation of individuals that will have relatives, and an independent set of individuals with no relatives.

Within each subsequent generation a constrained pairing of a male and female from the previous generation produced offspring. The pairings generated between individuals were constrained based on previous US census information (17). Within each generation, the probability that an individual would marry given they are male is:

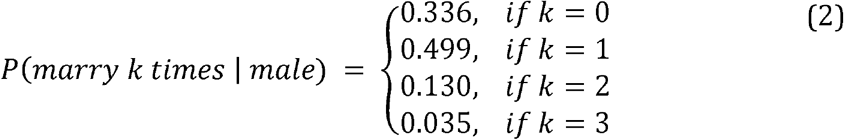

and the probability that an individual would marry given they are female is:

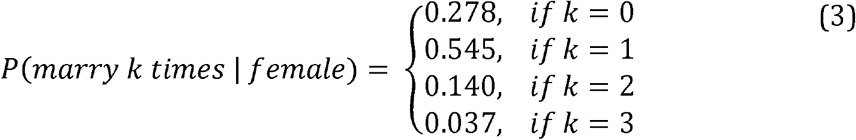

This information provides the probability that a person would get married, zero, one, two, or three or more times.

Furthermore, an ethnic intermarriage rate

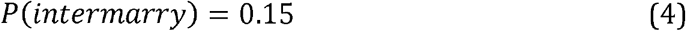

was set based on a poll conducted by pew social trends (18). If an individual were the product of intermarriage, they would be given a uniform probability of marrying all individuals of the opposite gender, regardless of their ethnicity. All couples that married produced four offspring. Each child was created independently with their two alleles drawn randomly from their parents at each locus. This procedure was followed for the remaining eight generations. This in silico profiles dataset is available (16).

### Characterizing Differences Between Ethnic Groups

The Euclidean distance was computed between minor allele frequencies across ethnic groups in order to better understand the distance between ethnic groups. This distance was represented as:

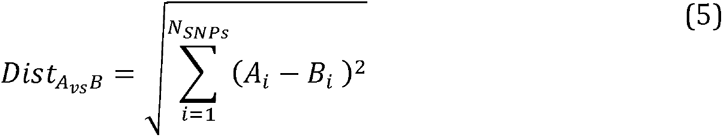

where A_i_ is the minor allele frequency for locus i within ethnicity A, and B_i_ is the minor allele frequency for locus i within ethnicity B.

Additionally, the fixation index (F_ST_) was computed as a standard method for identifying population differentiation due to genetic structure. F_ST_ was defined as the difference between the total population heterozygosity (H_T_) and the average subpopulation heterozygosity (H_s_), all divided by the total population heterozygosity. The formula for F_ST_ is shown in equation 6.

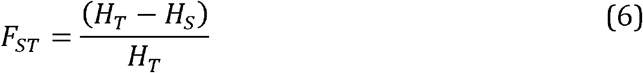

### Characterizing Differences Between Individuals

The differences between populations were also characterized using the mean number of mismatches between individuals. Across individuals, the number of mismatches is defined by the number of occurrences where an individual has at least one minor allele at a given locus, while this allele is absent in the second individual, or when the first individual is missing a minor allele that the second individual has at least one copy of. The case where one person has a single minor allele and the second person is homozygous for the minor allele is not counted as a mismatch. This is done to allow efficient querying of minor allele differences by making a binary decision based on whether or not there exists a minor allele at the given loci. All results are reported using two subsets of the panel of SNPs. The first subset contains 3,453 SNPs with a MAF between 0.01 and 0.1, while the second subset contains the 12,456 SNPs with a MAF between 0.01 and 0.2.

The predicted number of allele mismatches is derived using the Hardy-Weinberg equation:

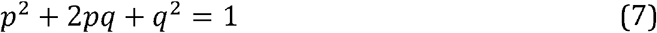

where p is the major allele frequency, and q is the minor allele frequency. Using this equation, the mean number of allele mismatches between unrelated individuals, defined here as the Combined Search Method, can be predicted as:

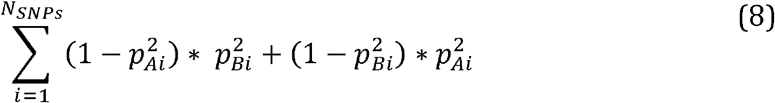

where P_Ai_ is the major allele frequency for population A, and P_Bi_ is the major allele frequency for population B, and 2*pq* + *q*^2^ = 1 — *p*^2^. This equation provides the sum of the probabilities that an individual from population A will have at least one minor allele, while an individual from population B will have two major alleles and vise versa. Furthermore, the number of differences at the loci where an individual has a minor allele and the second individual has only the major allele, defined here as the Individual Search Method, can be predicted as

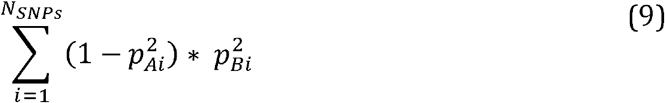

This equation provides a means for predicting the number of minor alleles that are present in an individual but absent from the population that they are being compared against.

### Admixtures and Kinship Mismatches

The framework for identifying minor allele mismatches can be extended to related individuals of both pure and mixed ancestries (admixture). A person with admixed ancestry has a combination of SNP frequencies drawn proportionally from minor allele frequencies of their inherited ancestries. The admixture SNP frequencies are represented by the following formulae:

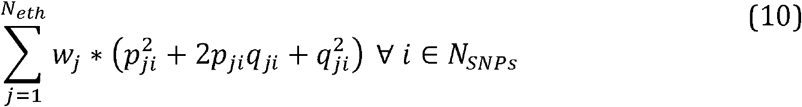

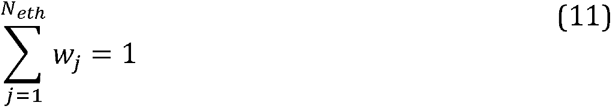

This allows for an individual to be compared against their descendants who have a mixed ethnicity. The prediction for expected number of minor allele mismatches once again builds on Hardy-Weinberg population genetics and probability theory. Equations were derived to predict the expected number of differences between parent and child, sibling and sibling, and grandparent and grandchild.

Given a child who is the product of a parent of ethnicity A, and a parent of ethnicity B, the expected number of differences between the child and the parent of ethnicity A can be computed in two stages. The first stage is to compute the probability that a child will have a number of minor alleles *n* ∈ {0,1,2}, given that parent A has a defined number of minor alleles *m* ∈ {0,1,2}.

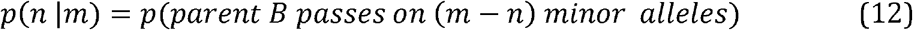

The second stage is to multiply by the prior probability that parent A will have a number of minor alleles, *m.*

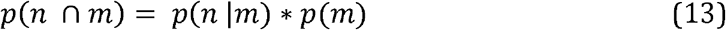

These relationships are computed and shown in Table 1. Additionally, the final joint probability is shown for the scenario where the parents have the same ethnicity (A = B)

**Table 1:**
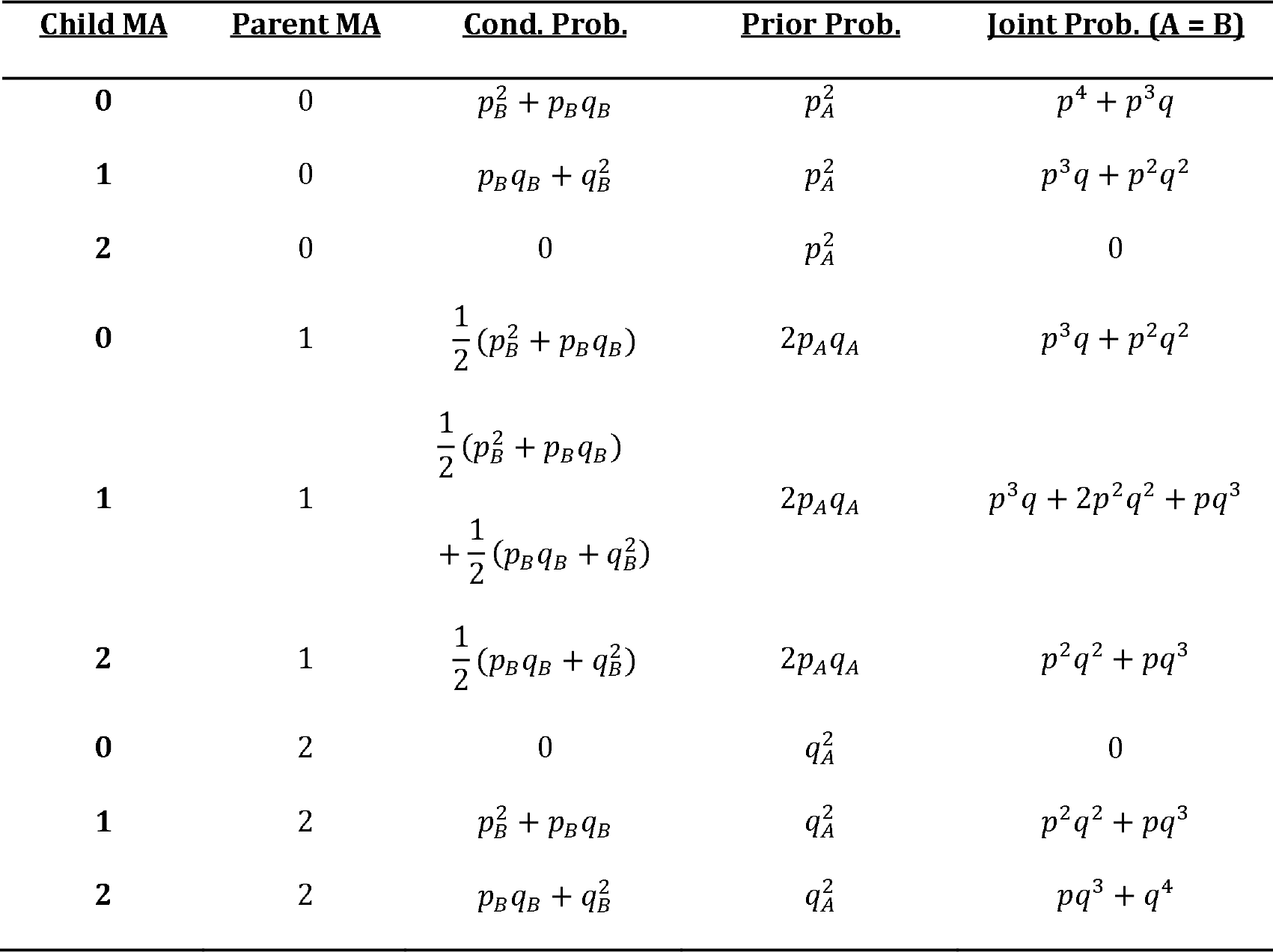
Table of probability equations used to estimate the number of mismatches between a child and their parent. The child is compared to a parent of ethnicity A, and has another parent of ethnicity B. The joint probability is shown for the condition when the parents are of the same ethnicity.

Using Table 2, it becomes possible to calculate the expected number of minor allele mismatches, which will be represented as the sum across loci of the probability that a child has 1 minor allele when their parent has 0 minor alleles, and the probability that the child has 0 minor alleles and their parent has 1 minor allele. In the event the parent and child are of the same ethnicity, the sum of the probabilities can be expressed as the equation:

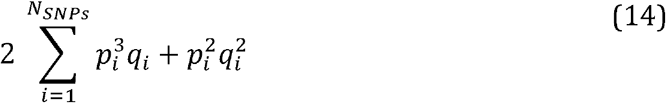

**Table 2:** Table of probability equations used to estimate the number of mismatches between an individual and their siblings. The siblings have two parents, one of ethnicity A, and the other of ethnicity B. The joint probability is shown for the condition when the parents are of the same ethnicity.

| <u>Sib1, Sib2</u> | <u>Parent1,</u><br><u>Parent2</u> | <u>Cond. Prob.</u> | <u>Prior Prob.</u> | <u>Joint Prob. (A = B)</u> |
| --- | --- | --- | --- | --- |
| 0, 1 | 0, 1 | $\frac{1}{4}$ | $p_A^2 * 2p_Bq_B$ | $\frac{1}{2}p^3q$ |
| 0, 1 | 1, 0 | $\frac{1}{4}$ | $p_B^2 * 2p_Aq_A$ | $\frac{1}{2}p^3q$ |
| 0, 1 | 1, 1 | $\frac{2}{16}$ | $2p_Aq_A * 2p_Bq_B$ | $\frac{1}{2}p^2q^2$ |
| 0, 2 | 1, 1 | $\frac{1}{16}$ | $2p_Aq_A * 2p_Bq_B$ | $\frac{1}{4}p^2q^2$ |

However, if the child has mixed ethnicity, with one parent of ethnicity A, and another parent of ethnicity B, then the sum of the differences will be expressed as the equation:

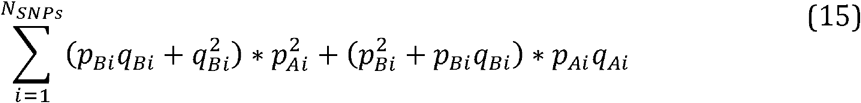

The framework for computing the expected number of minor allele mismatches also generalizes across the type and level of relationship. In order to compute the expected number of differences between two siblings, it is essential to account for the number of minor alleles each parent has at a given locus. Parent 1 is represented as being a member of ethnicity A, and parent 2 is represented as being a member of ethnicity B. The first equation computes the conditional probability of two siblings having a number of minor alleles *n*_1_*, n*_2_ ∈ {0, 1, 2}, conditioned on all the possible allelic combinations that their parents could have *m*_1_, *m*_2_ ∈ {0,1,2}. The conditional probability *p*(*n*_1_,*n*_2_ |*m*_1_,*m*_2_) is multiplied by the prior probability *p*(*m*_1,_*m*_2_), and is then marginalized over for all possible values of *m*_1_ and *m*_2_.

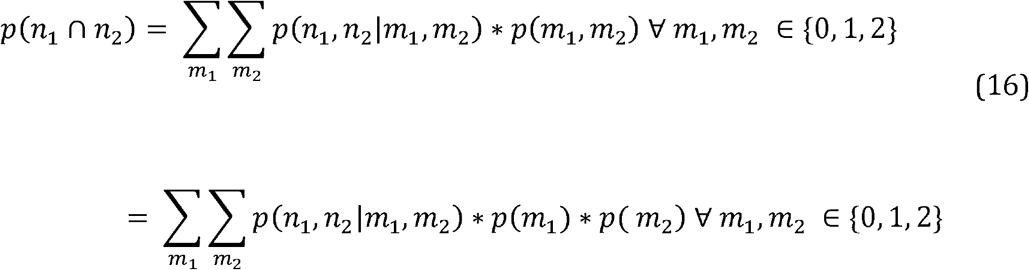

Because *m*_1_, and *m*_2_ are independent events, the joint probability *p* (*m*_1_, *m*_2_), can be represent as the product of *p* (*m*_1_) and *p* (*m*_2_). By marginalizing over the prior, the equation represents the full joint probability of differences between siblings. When examining differences between zero minor alleles and one or two minor alleles, the set of equations can be summarized as (zero probability parent child combinations not shown):

Using Table 2, two siblings of the same ethnicity will be represented by the equation:

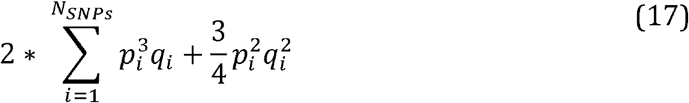

However, if the siblings have mixed ancestry, with one parent of ethnicity A, and the other of ethnicity B, the expected number of differences between siblings can be represented as:

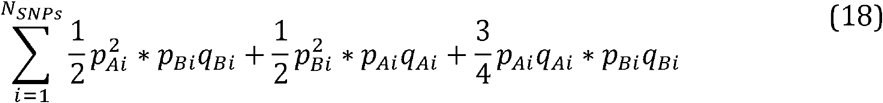

Finally, this methodology was employed to predict the expected number of minor allele mismatches between a grandparent and grandchild. This equation is derived using the chain rule of probability, alongside the conditional and prior probabilities computed in Table 1. Using the chain rule, the conditional probability is computed such that a child will have a number of minor alleles *n* ∈ {0,1,2}, given their grandparent has a number of minor alleles *ο* ∈ {0,1,2}, which factors into the sum of the product of conditional probabilities of a child having *n* minor alleles given their parent has *m* ∈ {0,1,2} minor alleles, and a parent having *m* minor alleles given their grandparent has *o* minor alleles. The summation of the conditional probabilities, multiplied by their priors is then used to compute the joint probability of a child having *n* minor alleles and their grandparent having *m* minor alleles. This results in four equations to compute the expected number of mismatches between a child and their grandparent.

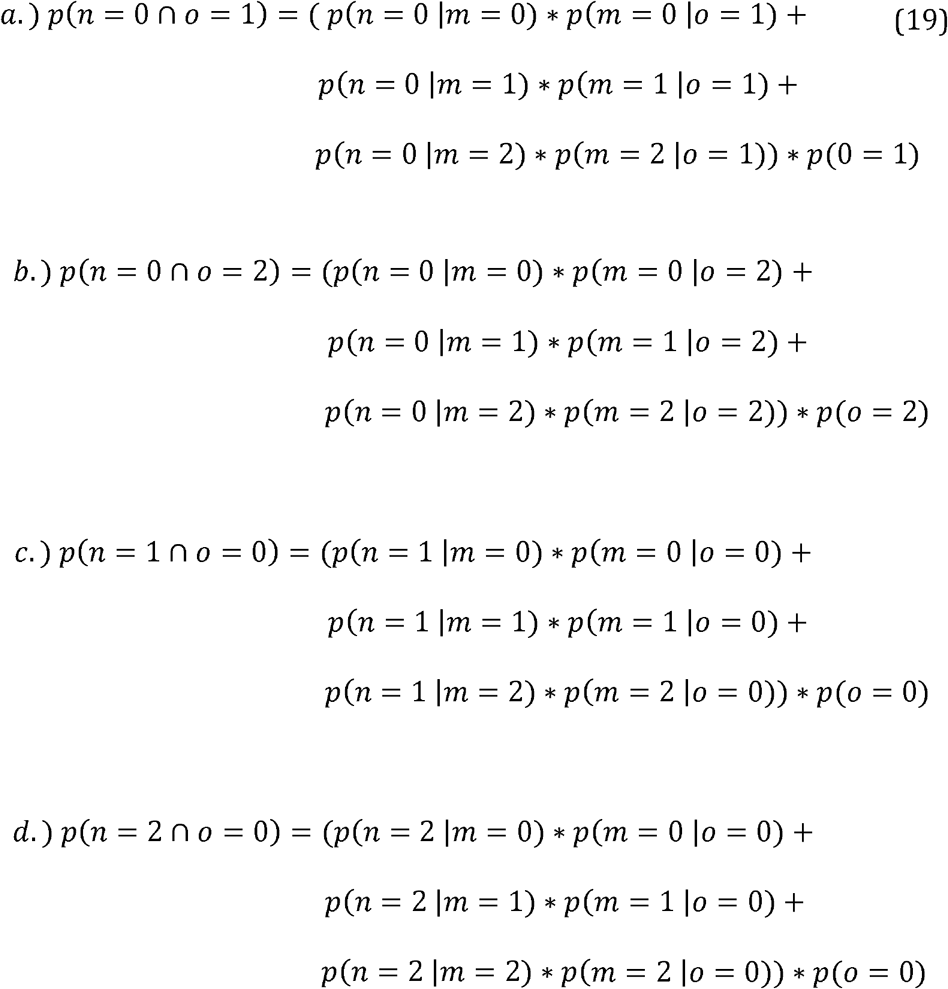

The summation of equations 19a through 19d over all SNPs produces the expected number of mismatches between a grandparent and grandchild.

### Kinship Equations for Computing Total Differences

The framework shown above can also be used to predict total mismatches between related individuals, where a mismatch is defined as a difference between genotypes. This differs from the above methodology, as it would add in a factor to represent mismatches where an individual is heterozygous for the minor allele, and their relative is homozygous for the minor allele. Using this definition, along with Table 1, the total expected differences between a parent and child of the same ethnicity would be:

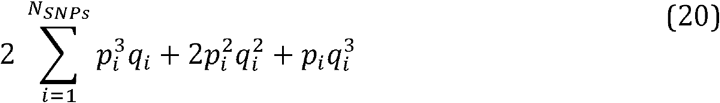

Similarly, the expected number of differences between siblings can be extended to account for the total number of differences between genotypes at the examined loci. Doing this requires the incorporation of the probability that an individual will have one minor allele, and their sibling will have two minor alleles. These probabilities are computed in Table 3.

**Table 3:** Table of probability equations used to estimate the number of genotype mismatches between an individual and their siblings. The siblings have two parents, one of ethnicity A, and the other of ethnicity B. The joint probability is shown for the condition when the parents are of the same ethnicity. This table extends upon Table 2.

| <u>Sib1, Sib2</u> | <u>Parent1,</u><br><u>Parent2</u> | <u>Cond. Prob.</u> | <u>Prior Prob.</u> | <u>Joint Prob. (A = B)</u> |
| --- | --- | --- | --- | --- |
| 1, 2 | 1, 1 | $\frac{2}{16}$ | $2p_Aq_A * 2p_Bq_B$ | $\frac{1}{2}p^2q^2$ |
| 1, 2 | 1, 2 | $\frac{1}{4}$ | $2p_Aq_A * q_B^2$ | $\frac{1}{2}pq^3$ |
| 1, 2 | 2, 1 | $\frac{1}{4}$ | $2p_Bq_B * q_A^2$ | $\frac{1}{2}pq^3$ |

If the siblings have two parents of the same ethnicity, then the expected number of total minor allele difference would be expressed as:

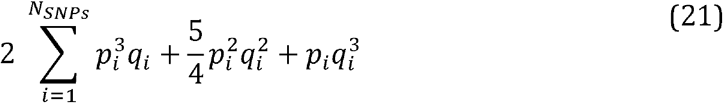

Finally, to predict the total expected number of differences between a grandparent and grandchild, two additional equations are incorporated.

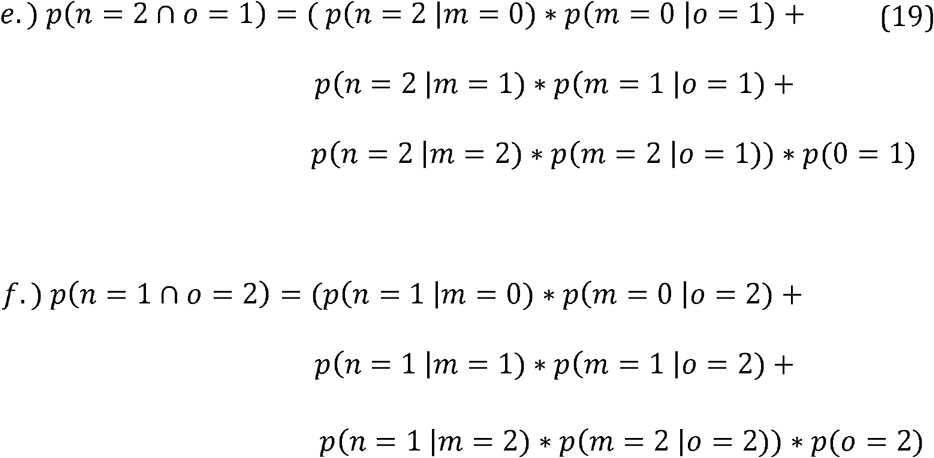

Factoring equations 19e and 19f into the summation over SNPs results in the prediction of the expected number of different genotypes across all loci.

## Results

Table 4 provides a summary of the Euclidean distance between each of the four ethnic groups. The Euclidean distance is computed to measure the similarity between each ethnic group as a function of the minor allele frequencies averaged over the population.

**Table 4:** Population distance metrics between each of the four ethnic groups a) Euclidean distance for the 3,453 SNPs with a MAF between 0.01 and 0.1. b) Euclidean distance for the 12,456 SNPs with a MAF between 0.01 and 0.2. c) F_ST_ for the 3,453 SNPs with a MAF between 0.01 and 0.1. d) F_ST_ for the 12,456 SNPs with a MAF between 0.01 and 0.2.

| **Ethnicity** | **African Am.** | **Estonian** | **Korean** | **Palestinian** |
| --- | --- | --- | --- | --- |
| **African Am.** | 0 | 5.48 | 5.62 | 5.17 |
| **Estonian** | 5.48 | 0 | 5.27 | 3.56 |
| **Korean** | 5.62 | 5.27 | 0 | 5.34 |
| **Palestinian** | 5.17 | 3.56 | 5.34 | 0 |

**(4b)**
| **Ethnicity** | **African Am.** | **Estonian** | **Korean** | **Palestinian** |
| --- | --- | --- | --- | --- |
| **African Am.** | 0 | 16.96 | 19.73 | 15.90 |
| **Estonian** | 16.96 | 0 | 15.48 | 8.52 |
| **Korean** | 19.73 | 15.48 | 0 | 16.05 |
| **Palestinian** | 15.90 | 8.52 | 16.05 | 0 |

**(4c)**
| **Ethnicity** | **African Am.** | **Estonian** | **Korean** | **Palestinian** |
| --- | --- | --- | --- | --- |
| **African Am.** | 0 | 0.0299 | 0.0358 | 0.0274 |
| **Estonian** | 0.0299 | 0 | 0.0302 | 0.0114 |
| **Korean** | 0.0358 | 0.0302 | 0 | 0.0311 |
| **Palestinian** | 0.0274 | 0.0114 | 0.0311 | 0 |

**(4d)**
| **Ethnicity** | **African Am.** | **Estonian** | **Korean** | **Palestinian** |
| --- | --- | --- | --- | --- |
| **African Am.** | 0 | 0.0458 | 0.0620 | 0.0410 |
| **Estonian** | 0.0458 | 0 | 0.0404 | 0.0122 |
| **Korean** | 0.0620 | 0.0404 | 0 | 0.0434 |
| **Palestinian** | 0.0410 | 0.0122 | 0.0434 | 0 |

**Table 5:**
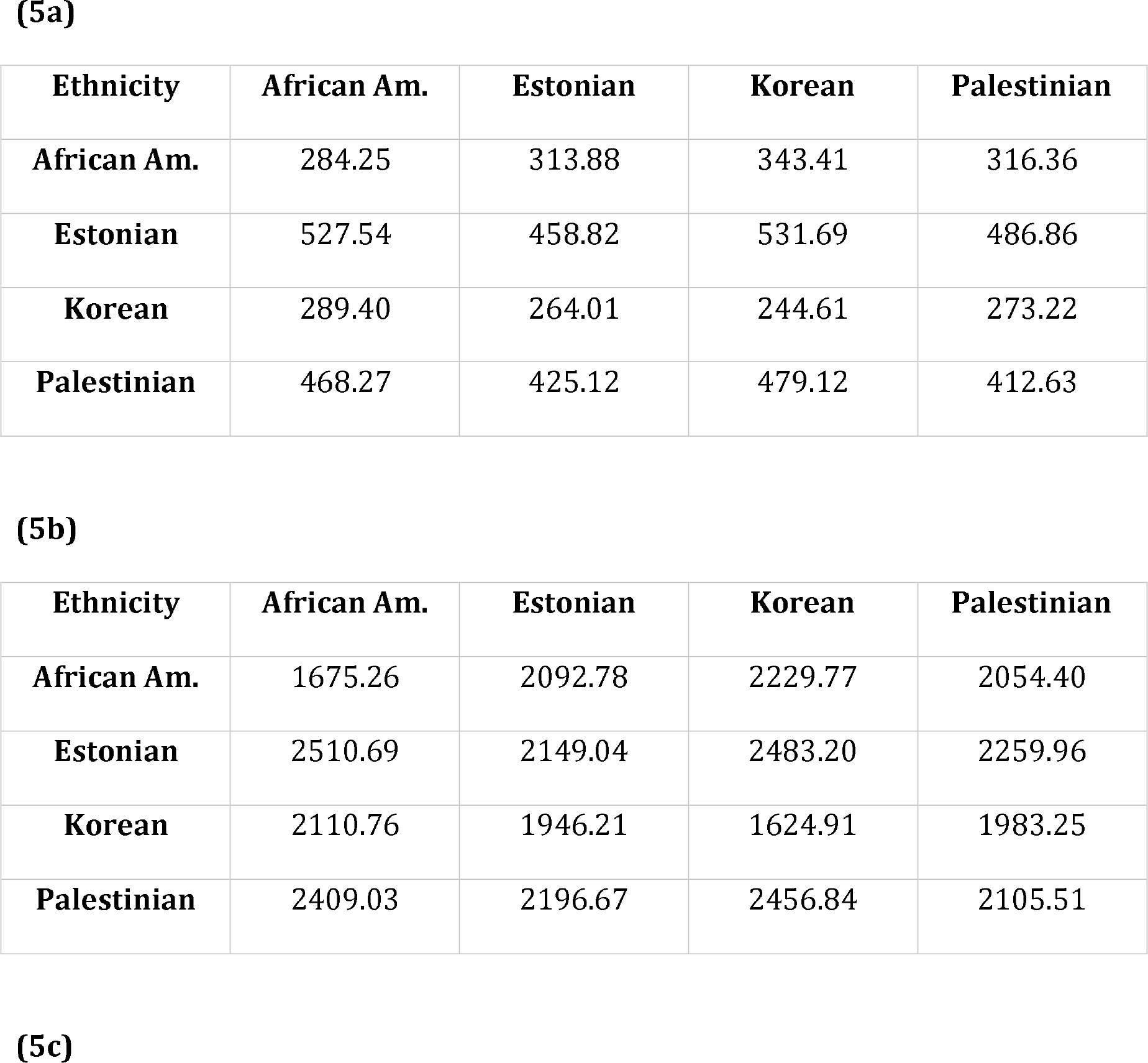
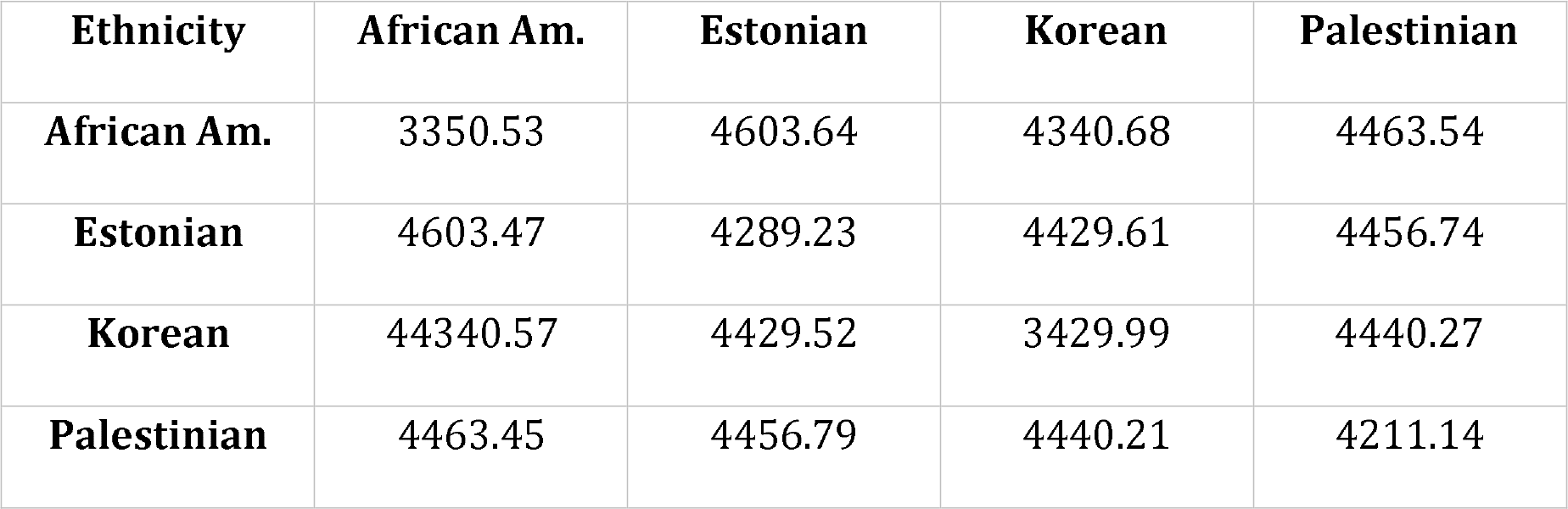
The population distance between each of the four ethnic groups a) Shown for the 3,453 SNPs with a MAF between 0.01 and 0.1 using the individual’s minor alleles. b) Shown for the 12,456 SNPs with a MAF between 0.01 and 0.2 using the individual’s minor alleles. c) Shown for the 12,456 SNPs with a MAF between 0.01 and 0.2 using all allelic differences.

While the Euclidean distance is informative when characterizing population differences, it is less effective when comparing differences between individuals. To meet this need, changes between ethnicities and individuals across multiple generations of the *in silico* database are tracked using the number of minor allele differences. This information is then plotted as a set of kernel density estimates (KDEs). These plots provide a distribution of the number of minor allele mismatches within each comparison. Figure 1 shows a set of KDE plots with the subset of loci with a MAF between 0.01 and 0.1 (Figure 1a) and the subset of loci with a MAF between 0.01 and 0.2 (Figure 1b). These minor allele frequency sets were selection for their potential use in DNA mixture analysis. The population of 15,000 individuals is compared to the set of unrelated individuals. The number of minor allele mismatches between groups is averaged across these comparisons.

**FIG 1.**
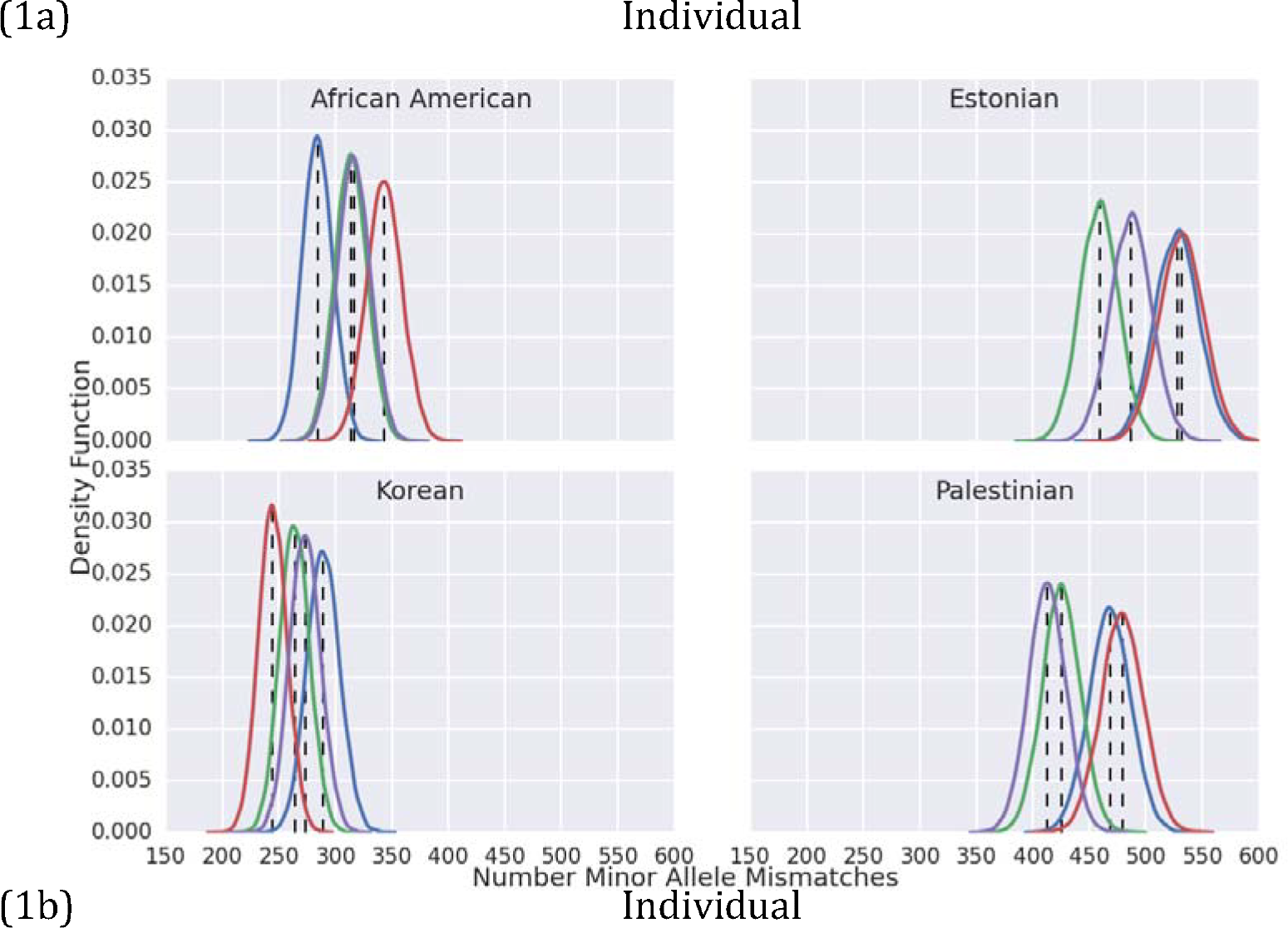
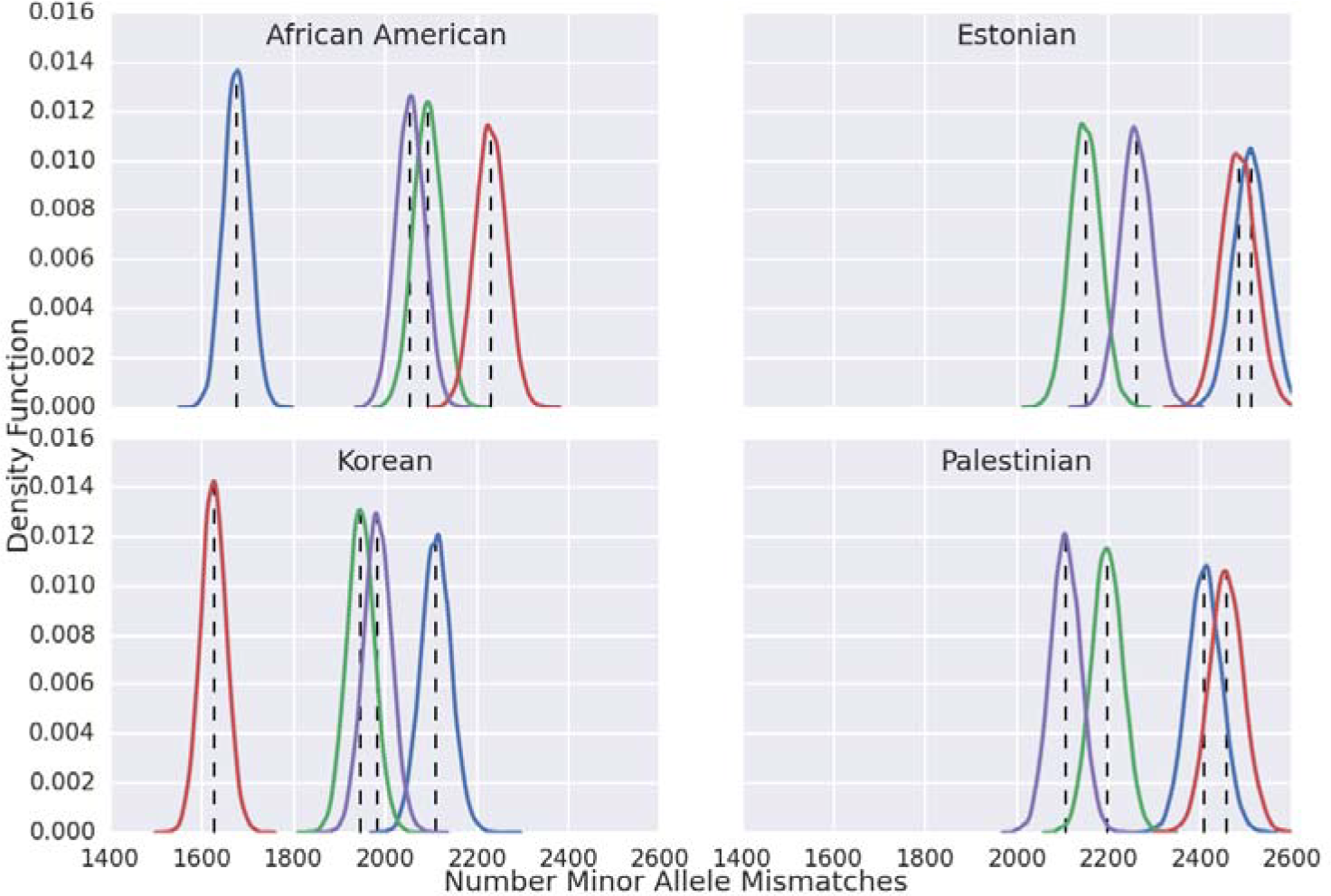
Mean Mismatch Individual Search Results: Blue = African American, Green = Estonian, Red = Korean, Purple = Palestinian, Black = Estimates. a) Shows the KDE for the mean comparison against the individual’s minor alleles with MAF between 0.01 and 0.1 (3,453 SNPs). b) Shows the KDE for the mean comparison against the individual’s minor alleles with MAF between 0.01 and 0.2 (12,456 SNPs).

Different methodologies are compared for identifying differences between individuals and ethnic groups. Figure 2 shows two different algorithms for identifying differences. The first algorithm, shown in Figure 2a, identifies the number of differences where an individual has a minor allele at a locus, while the person they are being compared to does not. The second algorithm, shown in Figure 2b, shows the total number of allelic differences.

**FIG 2.**
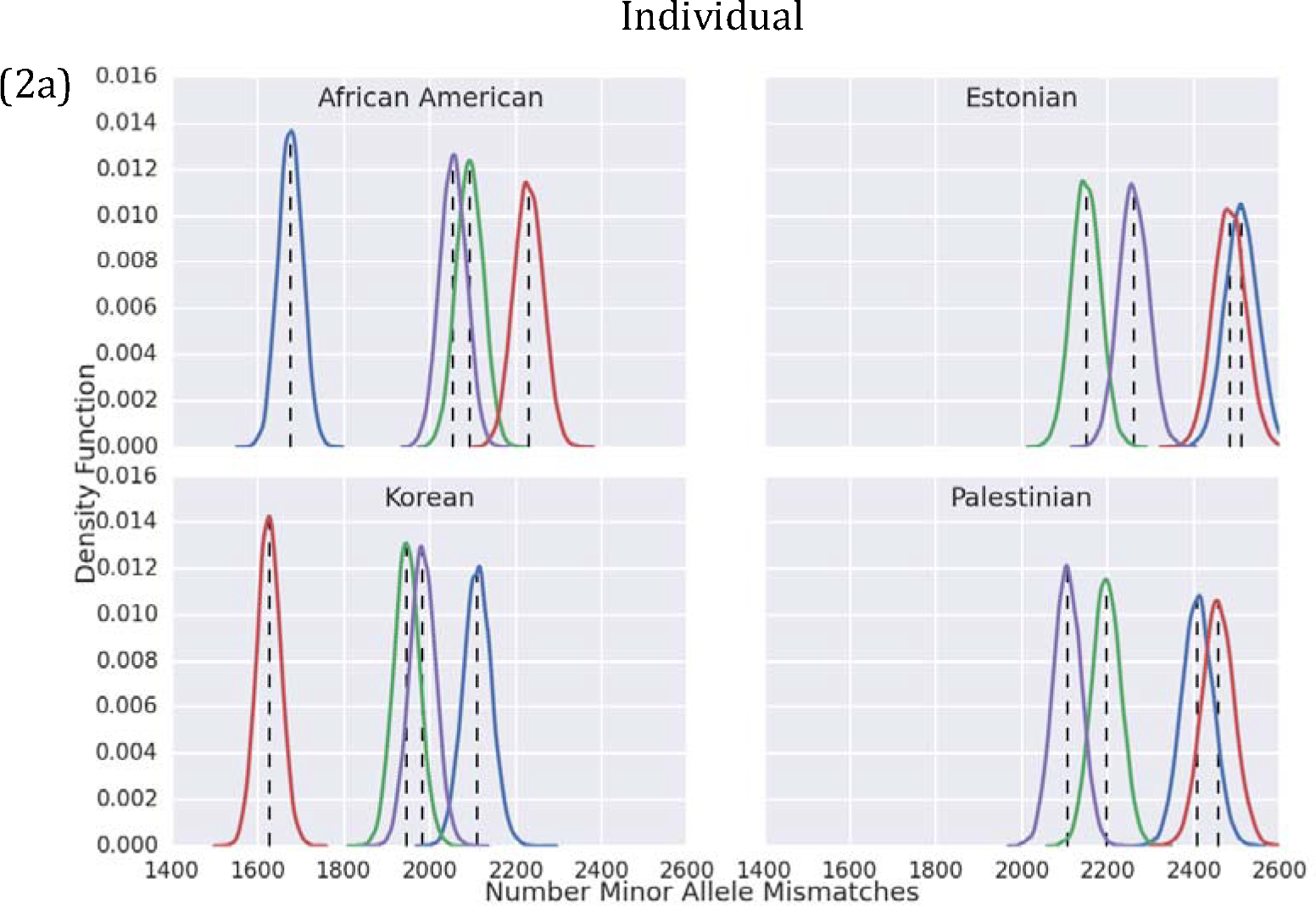
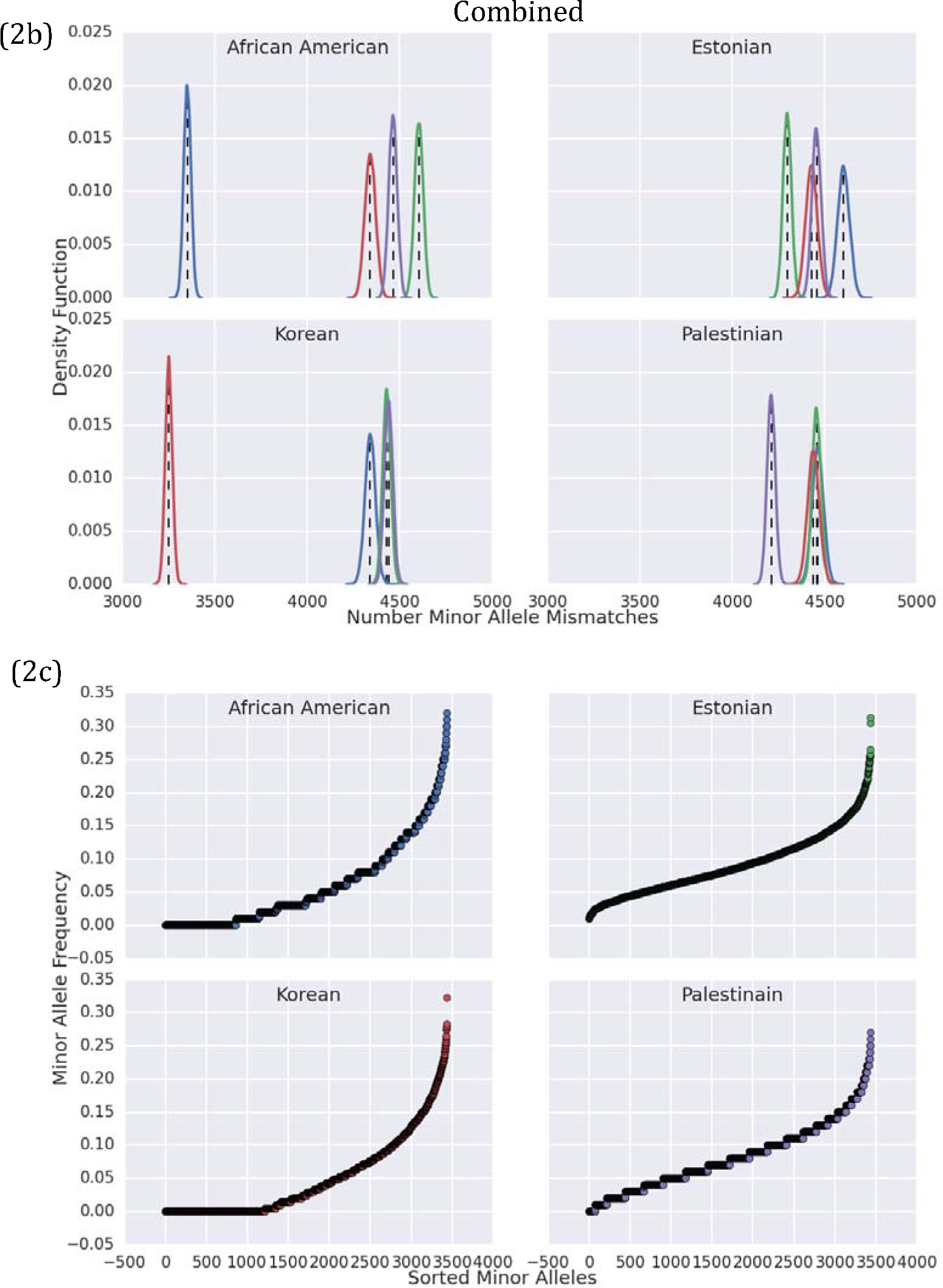
Mean Mismatch Individual and Combined Search Results: Blue = African American, Green = Estonian, Red = Korean, Purple = Palestinian. Black = Estimates. a) Shows the KDE for the mean comparison against the individual’s minor alleles. b) Shows the KDE for the mean comparison of all combined allelic differences. c) Shows the sorted MAF for each ethnic group.

Figure 2 shows an interesting phenomenon. Upon utilizing the set of all minor allele differences, as shown in Figure 2b, there is no longer a guarantee that the population of individuals will be most closely related to the ethnicity with the most similar profile of minor allele frequencies. For example, while Estonians and Palestinians are shown to be the most closely related ethnic groups (Table 4), in the Estonian and Palestinian quadrants of Figure 2b, we see that the Korean population is more closely related than the most similar ethnic group. This can be explained using the information shown in Figure 2c, which plots the sorted MAF across loci. From this plot it can be observed that the Korean population has approximately 1,200 loci with a MAF approaching zero. As a result of this, when all minor allele mismatches are tracked, as is the case in Figure 2b, the Korean population will show a smaller number of mismatches than more closely related foreign populations.

Figure 3 plots the regressed best-fit lines between Euclidean distances and population distances, showing high correlation between the two methods. In addition, a bootstrapping procedure is used to identify the 95% confidence intervals, which are shown as the shaded region around the blue curve.

**FIG 3.**
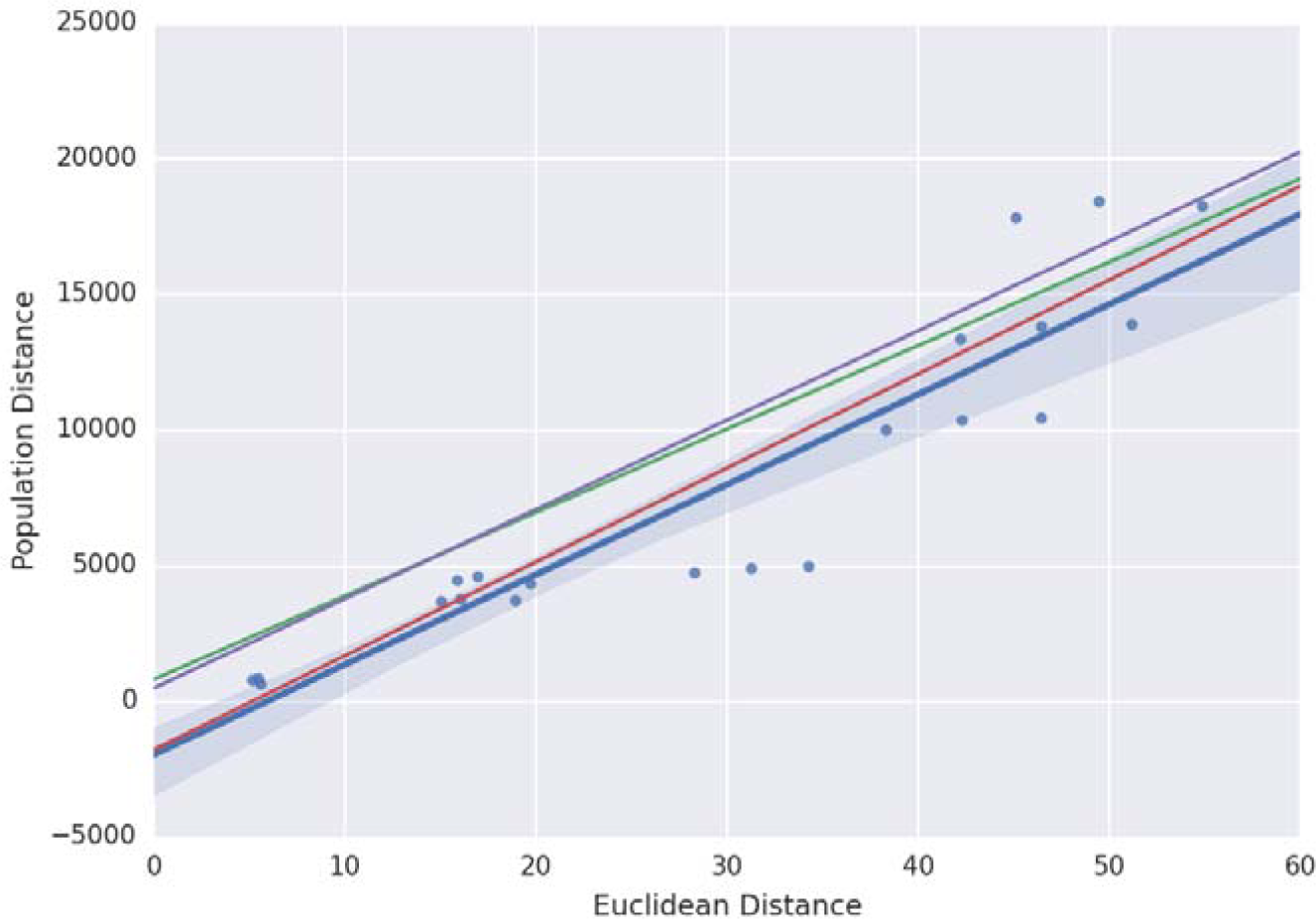
The four curves represent the correlation between Euclidean distances and population distances (that include all allelic differences) across the four ethnic groups: Blue = African American, Green = Estonian, Red = Korean, Purple = Palestinian. The African American correlation includes bootstrapped 95% confidence intervals.

The minimum number of minor allele mismatches across a population is also recorded as a way of assessing the lower limit of relation between differing populations. Figure 4 presents the minimum number of differences when comparing minor alleles found in the individual using the subset of loci with a MAF between 0.01 and 0.1, and the subset with a MAF between 0.01 and 0.2.

**FIG 4.**
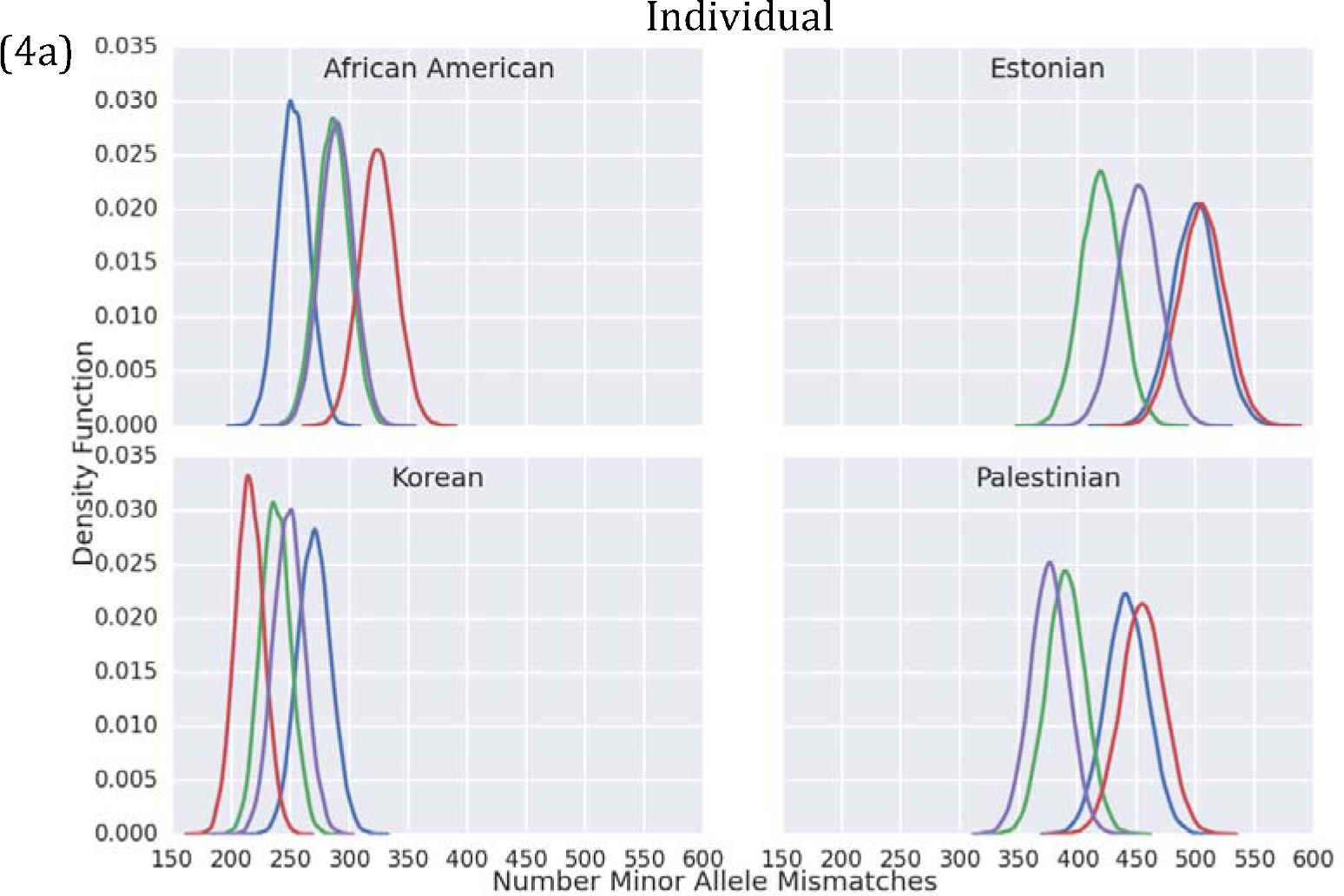
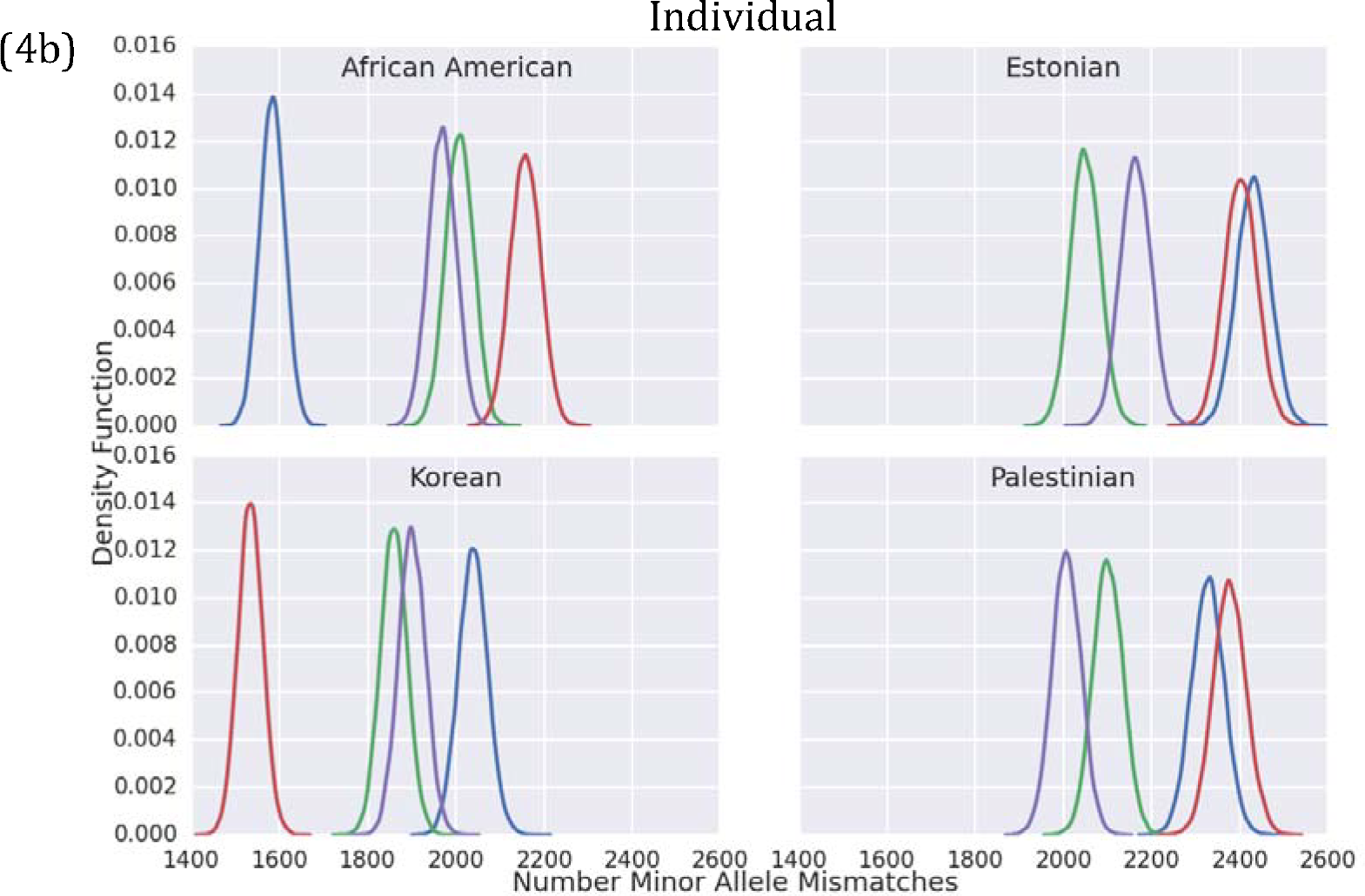
Minimum Mismatch Individual Search Results: Blue = African American, Green = Estonian, Red = Korean, Purple = Palestinian. Black = Estimates. a) Shows the KDE for the minimum comparison against the individual’s minor alleles with MAF between 0.01 and 0.1. b) Shows the KDE for the minimum comparison against the individual’s minor alleles with MAF between 0.01 and 0.2.

The *in silico* dataset allows for tracking changes in minor allele distribution across related individuals. This is demonstrated through a family that spans five generations. In each generation, some of the children married within their own ethnic background, while others married outside of their ethnic group. The observed intermarriage results in descendants with a varying percentage of their original ethnicity. This is demonstrated in Figure 5, which tracks an African American family across four generations. Each quadrant represents a degree of difference from the original ancestor (i.e. 1° = children, 2° = grandchildren, 3° = great grandchildren, 4° = great great grandchildren).

**FIG 5.**
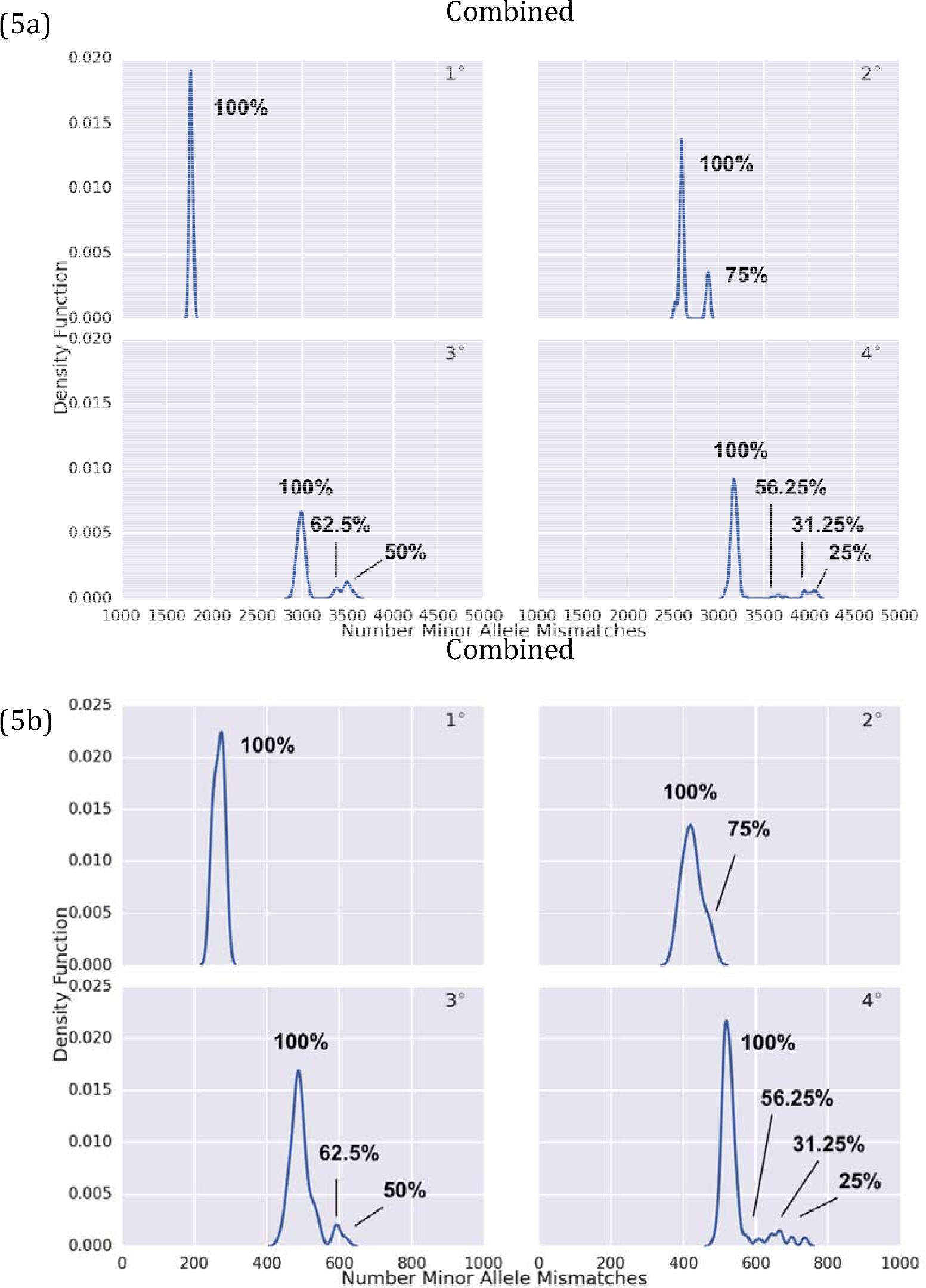
Results for close relatives including admixture. (Percentages identify the fraction of progenitor ethnicity shown in the distribution) a) Shows the KDE for 12,456 SNPs with a MAF between 0.01 and 0.2. b) Shows the KDE for 3,453 SNPs with a MAF between 0.01 and 0.1.

As can be observed from Figure 5, a greater dilution in the original ethnicity results in a greater number of mismatches across generations. This will result in someone with 75% of the original ethnicity with a second degree relationship, appearing similar to someone with 100% of the parent ethnicity and a third degree relationship. In order to capture this affect across families, mismatches between thousands of descendants was tracked at the parent-child, and grandparent-grandchild level. Figure 6 shows the average level of parent-child, and grandparent-grandchild relation, when the descendant has 100%, 75%, 50%, and 25% of the ethnicity of the progenitor. Each descendant has at most two races that go into their ethnic background.

**FIG 6.**
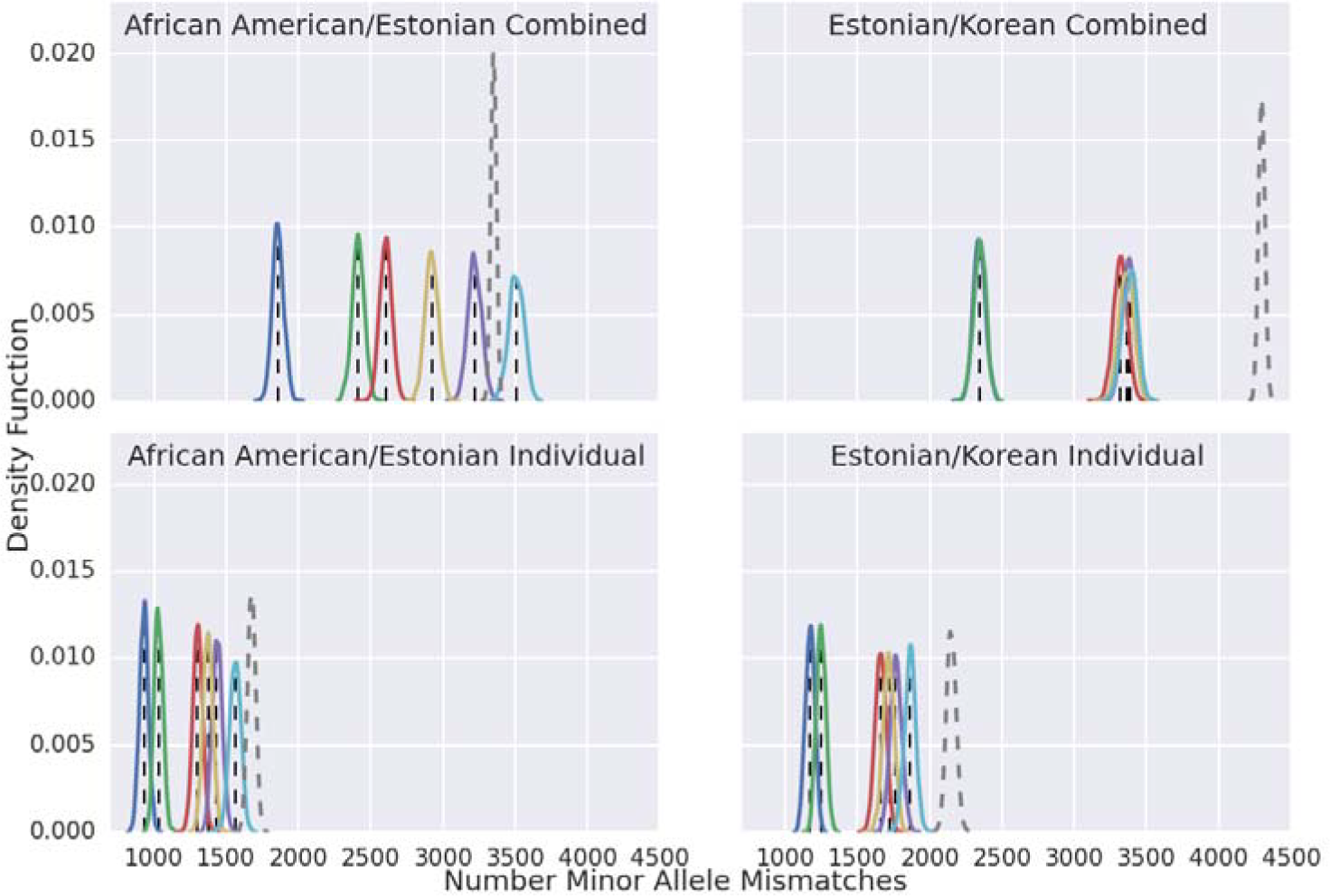
Kinship generations with admixture. (Blue = Gen2,100% progenitor race; Green= Gen2, 50% progenitor race; Red = Gen3,100% progenitor race; Yellow = Gen3, 75% progenitor race; Purple = Gen3, 50% progenitor race; Cyan = Gen3, 25% progenitor race; Gray = Unrelated progenitor race; Black dash = Predictions based on probability equations)

## Discussion

The probability equations used to estimate the number of mismatches between related and unrelated individuals builds on the work of Cassa *et al* (18). However, while their work looks to identify how an individual’s genotype can be inferred from their sibling’s genotype, this work builds out the relationship between individuals of different levels of relationship, and then compares their expected values. This paper provides an in-depth analysis of changes in minor allele representation across separate ethnic groups as well as through generations of genetic inheritance.

Table 4 provides a summary of the Euclidean distance between each of the four ethnic groups. This provides a theoretical basis for interpreting the *in silico* results. Each ethnic group will be closest to their own ethnicity, followed by each of the remaining groups. Based on Table 4, African Americans are predicted to be most similar to Palestinians, followed by Estonians, and finally Koreans. Estonians will have the most in common with Palestinians, then Koreans, and finally African Americans. Koreans are closest to Estonians, followed by Palestinians, and then African Americans. Finally, Palestinians are closest to Estonians, followed by African Americans, and finally Koreans. The Euclidean distances also show that the shortest allelic distance between differing ethnicities is between Estonians and Palestinians. This information is concordant with the F_ST_ numbers that have also been shown in Table 4. Additionally, Figure 3 demonstrates that there is a strong correlation with narrow error bars between the Euclidean distances and the population distances. This creates a method for identifying similarity between ethnic groups, which then provides information on the expected distance between individuals in two different populations.

The curves shown in Figure 1a and Figure 1b demonstrate the increased ability to separate populations of individuals as a function of the number of incorporated SNPs. Figure 1b shows that the increased subset of SNPs provides a greater degree of separation between ethnic groups. Upon examining this plot, it can be observed that the order of closest ethnicity matches what is shown in Table 4 by Euclidean distances. The only exception to this is shown in the panel of Estonian plots where the African American curve is closer than the Korean curve. However, in this case the African American and Korean curves are significantly overlapping. Figure 2a and Figure 2b show a comparison of the Individual and Combined methods for identifying minor allele differences between individuals. The combined method provides for symmetry of comparisons, while the individual methods tailors the comparison to the minor alleles present in the person of interest. As shown in the combined method, this property of symmetry can cause a group of unrelated individuals from a particular ethnic group to have a distribution that is consistently closer to the individual’s ethnicity. This is a function of the number of loci with a minor allele frequency of approximately zero, as shown in Figure 2c. This large number of alleles with a near zero MAF cause a greater prevalence of the major allele, leading to less allelic mismatches with the combined method. This result causes the Korean population to be consistently closer in distribution to the other ethnic groups.

The comparative methods are extended to kinship generations with admixture. As shown in Figure 6, these methods can be employed to identify related individuals of mixed race. However, as the fraction of the progenitor’s original ancestry becomes minimized, it becomes more difficult to separate out related individuals from unrelated individuals of the same ethnicity.

## Conclusions

This paper has presented a framework for characterizing identity search results, which has been used to compare both related and unrelated individuals from four different ethnicities. In the process of distinguishing populations of different ethnicities, this work has shown that ethnicity impacts the expected number of minor allele mismatches as expected. Furthermore, this method has been applied to related individuals, where it has been shown that relatives with admixture ancestry have more minor allele mismatches than relatives with shared ancestry.

## Acknowledgements

Distribution Statement A. Approved for public release: distribution unlimited. This material is sponsored by the Defense Threat Reduction Agency under Air Force Contract No. FA8702-15-D-0001. Any opinions, findings, conclusions or recommendations expressed in this material are those of the author(s) and do not necessarily reflect the views of the U.S. Air Force.

